## Supplementary Information for "New Asgard archaea capable of anaerobic hydrocarbon cycling"

### Supplementary Materials

#### Supplementary Methods:

#### JGI generation of reads and processing of data

DNA was extracted from sediment samples using the MO BIO – PowerMax Soil DNA Isolation kit[1]. A half a lane of Illumina reads (HiSeq–2500 1TB, read length of 2x151 bp) were generated at Joint Genome Institute for each sample, producing a total of 226,647,966 and 241,605,888 reads from dives for 4569-2 and 4571-4, respectively. The average percent of reads with a phred-score (Q) ≥ 30 was 86.2% and 90.39% and the average base quality score was 34.35 ± 7.73 and 35.38 ± 6.52 for samples from dive 4569-2 and 4571-4, respectively. The JGI performed read quality checks and generated a first assembly using the following methods: BBDuk adapter trimming removed known Illumina adapters. The reads were further processed using BBDuk quality filtering and trimming to remove reads quality score less than 12, containing more than three 'Ns', or with quality scores (before trimming) averaging less than 3 over the read length, or length under 51 bp after trimming. Additionally, reads matching Illumina artifacts or phiX were discarded. The remaining reads were mapped to a masked version of the human HG19 with BBMap and all hits over 93% sequence identity to the human genome were discarded. Trimmed, screened, paired-end Illumina reads were assembled using the megahit assembler using a range of kmers. Assemblies were preformed with default parameters in megahit with the following options: "--k-list 23,43,63,83,103,123". High-quality reads were mapped to the final assembly to calculate coverage information using bbmap by excluding all parameters except ambiguous=random as described by JGI.

#### Genome reconstruction and read abundance

The scaffolds from the assembly were binned based on tetranucleotide signatures and clustered using emergent self-organizing maps (ESOM[2], MetaBAT[3] and CONCOCT[4]. For ESOM, binning was performed on contigs with a minimum length of 2,000 bp using the K-batch algorithm for training after running the perl script esomWrapper.pl[2]. Emerging Self-Organizing Maps (ESOM) were manually sorted and curated. The bins were extracted using getClassFasta.pl (using -loyal 51). Reference genomes were included to add genetic signatures for the assembled contigs and improve binning. For CONCOCT, Anvi'o (v2.2.2) was used as the metagenomic workflow pipeline[5]. Coverage information was obtained by mapping all high-quality reads of each sample against the assembly of another sample using the BWA-MEM algorithm in paired-end mode (bwa-0.7.12-r1034; using default settings)[6]. The resulting sam file was sorted and converted to bam using samtools (version 0.1.19)[7]. The bam file was prepared for Anvi'o using the script anvi-init-bam and a contigs database generated using anvi-gen-contigs-database. These files were the input for anvi-profile. Generated profiles for the assemblies were combined using anvi-merge and the resulting bins summarized using anvi-summarize (-C CONCOT)[5]. If not mentioned otherwise, the scripts were used with default settings. Metabat was also used as a binning approach (v1)[3]. As described for Anvi'o the input consisted of the scaffold files (≥ 2000 bp) and the mapping files. First, each of the mapping files were summarized using jgi_summarize_bam_contig_depths and then metabat was run using the following settings: --minProb 75 --minContig 2000 --minContigByCorr 2000. Results from the three different binning tools were combined using DAS Tool (version 1.0)[8]. For each of the binning tools a scaffold-to-bin list was prepared and DAS Tool run on each of the eleven scaffold files as follows: DAS_Tool.sh -i Anvio_contig_list.tsv,Metabat_contig_list.tsv,ESOM_contig_list.tsv -l Anvio,Metabat,ESOM -c scaffolds.fasta --write_bins 1.

To improve the quality of the two Helarchaeota genomes IDBA-UD was run on raw data using the command: "idba_ud -r Guay9_METAGENOME.fasta -o G9 --pre_correction --mink 75 --maxk 105 --seed_kmer 55 --num_threads 30". Metaspades was run on Raw data and Metabat assembled bins using as follows: "metaspades.py --12 Guay16.11400.5.204846.CTCTCTA-CGTCTAA.filter-METAGENOME.fastq -o Metaspades --only-assembler --meta". Binning procedures described above were repeated with these new assembles. Final bins were selected from Metaspades assemblies.

Read abundance summarized by jgi_summarize_bam_contig_depths were used to calculate relative read abundance and total percent of metagenomic reads. Relative read abundance was calculated as total read abundance normalized to genome size and divided by total reads. Relative read abundance was then multiplied by the constant $1x10^{12}$ for clarity. Total percent of metagenomic reads was calculated as total read abundance divided by total reads times 100. Relative read abundance was compared to other genomics bins recovered from these sites to look for co-occurrence[9].

**Hydrogenase Analyses**

All hydrogenases detected in the bins were used to generate two phylogenetic trees, one for proteins identified as small subunits and one for large subunits in order to properly identify the different hydrogenase subgroups (Supplementary Figure 4). A small subunit tree was generated by aligning sequences from Helarchaoeta and published sequences as classified by Vignias et al.[10,11]in Geneious[12] and running a maximum-likelihood phylogenetic tree using the command raxmlHPC-PTHREADS-AVX -T 10 -f a -m PROTGAMMAAUTO -N autoMRE -p 12345 -x 12345 -s Protein_alignment_2_masked.phy -n tree_2. The large subunit tree was made using an alignment of sequences from HydDB as previously described[13] and the command raxmlHPC-PTHREADS-AVX -T 20 -f a -m PROTGAMMAAUTO -# 100 -p 12345 -x 12345 -s Protein_alignment_anja_edit.phy -n hydrogenase_anja_tree_2.

Included in the hydrogenase tree are three sequences from both bins that we proposed to be acting as a single complex. The HydB/Nqo4-like is affiliated with group III B (large subunit) and the Oxidored_q6 superfamily protein appears to be group IV hydrogenase-associated, which has been described to play a role in hydrogen trans-inner membrane exchange in other organisms30,31 (Supplementary Figure 4). Interestingly, while the Fe-S disulfide reductase/FlpD component falls within the small subunit of Group III hydrogenases, it does not appear to belong to any described subgroups suggesting that, despite its annotation, may not be mvh (Supplementary Figure 4). Nucleotide sequences were analyzed in Geneious[12] to evaluate if this was a possible operon by looking for possible transcription factors and binding motifs. Possible TATA boxes and BRE sequences were located on both contigs. The prodigal protein predictions were used to determine directionality and length of the potential operon. No consistent annotation was found for these three genes in either bin, however, a NCBI blastp search identified these hits as possible hydrogenases belonging to the superfamilies HydB-Nqo4 for NODE_1033_length_35804_cov_14.1912_29 and NODE_147_length_7209_cov_4.62199_7, oxidored_q6 for NODE_1033_length_35804_cov_14.1912_30 and NODE_147_length_7209_cov_4.62199_6, and HrdA/FlpD for NODE_1033_length_35804_cov_14.1912_31 and NODE_147_length_7209_cov_4.62199_5 (Figure 5a). These superfamilies were then confirmed using the online InterproScan system[14]. The TMHMM webserver[15], PRED_TMR webserver[16] and Phobius[17] was used to identify membrane motifs and positions relative to the membrane. Proteins were analyzed both individually and concatenated into a single sequence but no difference was seen between these two methods. All genes showed some membrane association, however, for the HydB-Nqo4-like protein identified in Hel_GB_A this may be the result of ambiguous bases in the sequence as removal of said bases removed the membrane signature. Results from all the transmembrane predictor programs were compared and consistent residues were found across all three programs with the exception of the second transmembrane region associated with the oxidored_q6 protein (Hel_GB_B AA918-934), which could not be found in PRED-TMR. Given the forward direction of the operon and the more reliable HydB-Nqo4 signature, Hel_GB_B was used to create a possible diagram of the amino acid orientation across the membrane (Figure 5b).

**ESP Identification**

The DNA polymerase epsilon subunit was considered absent when the specific IPR domain signature was not identified, even if the corresponding arCOG was detected. Regarding topoisomerase IB, BLAST verifications were carried out when the arCOG was not detected.

Helarchaeota have a fused version of RNA polymerase A, similarly to Heimdall_LC3 and eukaryotes. Phylogenetic analysis using all the available Asgard genomes was used to determine the origin of these fusions/splits. The alignment was generated using Muscle[18] and trimmed used BMGE (-m BLOSUM30)[19]. IqTree[20] was run using the best fit model (LG+R9). The tree was poorly resolved, but the Asgard sequences were monophyletic and, overall, the fused genes seemed to cluster according to the Asgard phylogeny (Helarchaeota with Lokiarchaeota). This suggests that multiple fusion and split events have happened in the evolution of this protein family in archaea. The eukaryotes do not branch with the Asgard homologs in this phylogeny.

The arCOG04271, which comprises homologues of the RNA polymerase subunit RPB8, was found in the Helarchaeota and confirmed by pfam[21] and Hhpred[22]. The Interpro IPR002671 family that contains the ribosomal protein L22e was detected in Helarchaeota, although their arCOG was different. SMART[23] finds a PF01776 (L22e) domain but with a very low e-value (0.001). IPR029004 (Ribosomal protein L28e) was not detected at all.

An ESCRT-I (Vps28) homologue belonging to IPR07143 was detected in Hel_GB_B but not in Hel_GB_A. However, when using the Vps28 homolog of Hel_GB_B (Hel_GB_B_02460), Hel_GB_A_08820 was detected with BLAST; no IPRs, arCOG or pfam profiles are detected in Hel_GB_A_08820. ESCRT-I steadiness box domain's IPR017916 domain, however, is found in both Hel_GB bins. For ESCRT-II Vps22/36 (EAP30), IPR007286 is present in Hel_GB_B but not Hel_GB_A. This was confirmed with BLAST. ESCRT-II Vps25 are found in two InterPro profiles, IPR014041 and IPR008570, which are both detected in Hel_GB_B, but not in Hel_GB_A, as subsequently confirmed by BLAST. Regarding ESCRT-III Vps2/24/46 and Vps20/32/60, IPR005024 is detected twice in Hel_GB_B but only once in Hel_GB_A (confirmed by BLAST). A phylogeny revealed that the unique Hel_GB_A copy belongs to the Vps20/32/60 group, which means that the Vps2/24/46 orthologue is lacking. Longins' IPR011012 and IPR004353 are present in both Hel bins.

In order to determine whether the Helarchaeota bins encode roadblock homologs that belong to the particular eukaryotic roadblock family (RLC7)[24], we performed phylogenetic analyses of the IPR004942/IPR015019 family proteins. Sequences from Helarchaeota were added to the alignment used in Zaremba-Niedzwiedzka, *et al*. 2017 using both Muscle[18] and Mafft[25]. Trimming was done with trimal[26] using the gappyout option and the phylogenetic reconstructions were made with FastTree[27]. In these phylogenies only one Hel_GB_A sequence clearly branches within the previously reported Asgard archaeal RLC7 family proteins and their close eukaryotic homologs.

TRAPP protein profiles IPR007194 and IPR024096 were not found in Hel_GB bins. This was confirmed by BLAST. As for Sec23/Sec24, the signature profiles IPR006895, IPR006896 and IPR012990 were not found. Two proteins containing IPR004000 were detected in both Helarchaeota, that represent conserved lokiactin homologues. Based on the detection of the InterPro profiles, both Helarchaeota should contain at least one additional actin-related protein belonging to IPR20902 (this was also confirmed in a phylogeny that contained homologs of all the Asgard genomes) but none related to IPR008384. BLAST supports the latter observation. Profilin's IPR005455 was detected in both genomes. For Gelsolin, we looked at IPR007122, IPR007123,

IPR029006, IPR029006, IPR029919, IPR030004 and arCOG20384. The arCOG was not detected and neither was the main IPR (IPR029919). The other IPRs, however, were detected in both genomes and confirmed with BLAST. Thus, we consider that these are good homologue candidates, although they may not be orthologues. IPR000217, IPR002453 and IPR023123, which are the specific IPRs for beta tubulins, were not detected in the new genomes.

IPR029071 and IPR000626, which are ubiquitin-domain proteins, were detected in both genomes. The RWD domain IPR006575 was detected in the Helarchaeota bins and was confirmed using BLAST. The profile of the ubiquitin-activating enzyme E1 (IPR000594) was found in both genomes, but its lack of specificity required a validation of the putative homology. The best markers of ubiquitin clusters are IPR019572 and IPR014929. Neither of these genomes carries IPR019572, whereas IPR014929 is only present in Hel_GB_A. We checked the synteny of IPR000594, IPR019572 and other ubiquitin-related genes. For instance, IPR000608 (putative E2-like protein), IPR013083 (putative E3-like proteins) and IPR018611 (UFM1-domain protein) were detected in a number of proteins in both genomes. Similarly, IPR000555 but not IPR028090 (both, putative deubiquitinating enzymes) homologues were detected in the Helarchaeota genomes. The synteny search showed that Hel_GB_A has a cluster that contains the IPR014929-bearing gene and other ubiquitin-related genes, but no genes containing the more specific IPR000594. Given these inconclusive results, we reconstructed a phylogeny of the proteins that carry an IPR000594 domain, based on the original alignment from Zaremba-Niedzwiedzka, *et al*. 2017. The Helarchaeota sequences were added using Mafft[25] and the alignment was trimmed with Trimal[26] (gappyout option). The resulting tree, which was generated using FastTree[27], revealed that the Helarchaeota sequences branch together with other Asgard ubiquitin candidates and close to the eukaryotic homologues. The sum of these analyses suggests that Hel_GB_A has good candidates of these proteins, whereas the Hel_GB_B has some candidates yet lacks other specific ubiquitin proteins.

Finally, ribophorin I (IPR007676) and STT3 subunit (IPR003674) homologues were detected in Hel_GB_B, but not in Hel_GB_A, as confirmed by BLAST. The fact that OST3/OST6 homologues (IPR021149) were detected in both Hels suggests that lack of a STT3 homologue in Hel_GB_A may be due to the incompleteness of the genome.

**Supplementary Figures:**

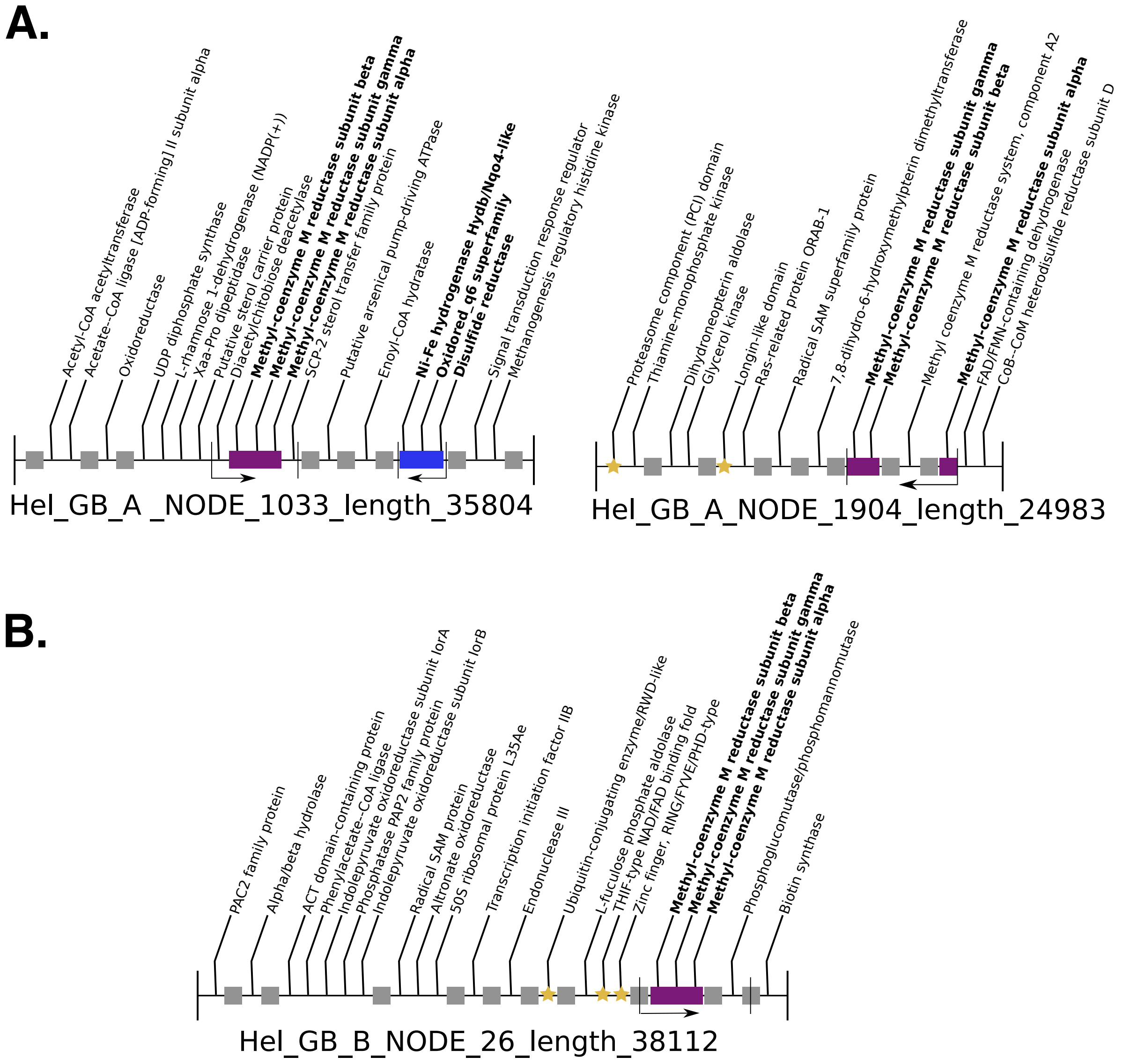


**Supplementary Figure 1.** Gene-neighborhood of contigs containing the methyl-CoM reductase operon in Helarchaeota. Hel_GB_A contained two mcrABC-containing contigs (A), while only one was found in Hel_GB_B (B). Purple blocks represent the mcrABC operon. Blue block represents possible electron transporting contig. Gold stars represent ESPs identified on the contigs and grey boxes symbolize hypothetical proteins. Arrows show the predicted directionality of the reading frame as predicted by prodigal[28].

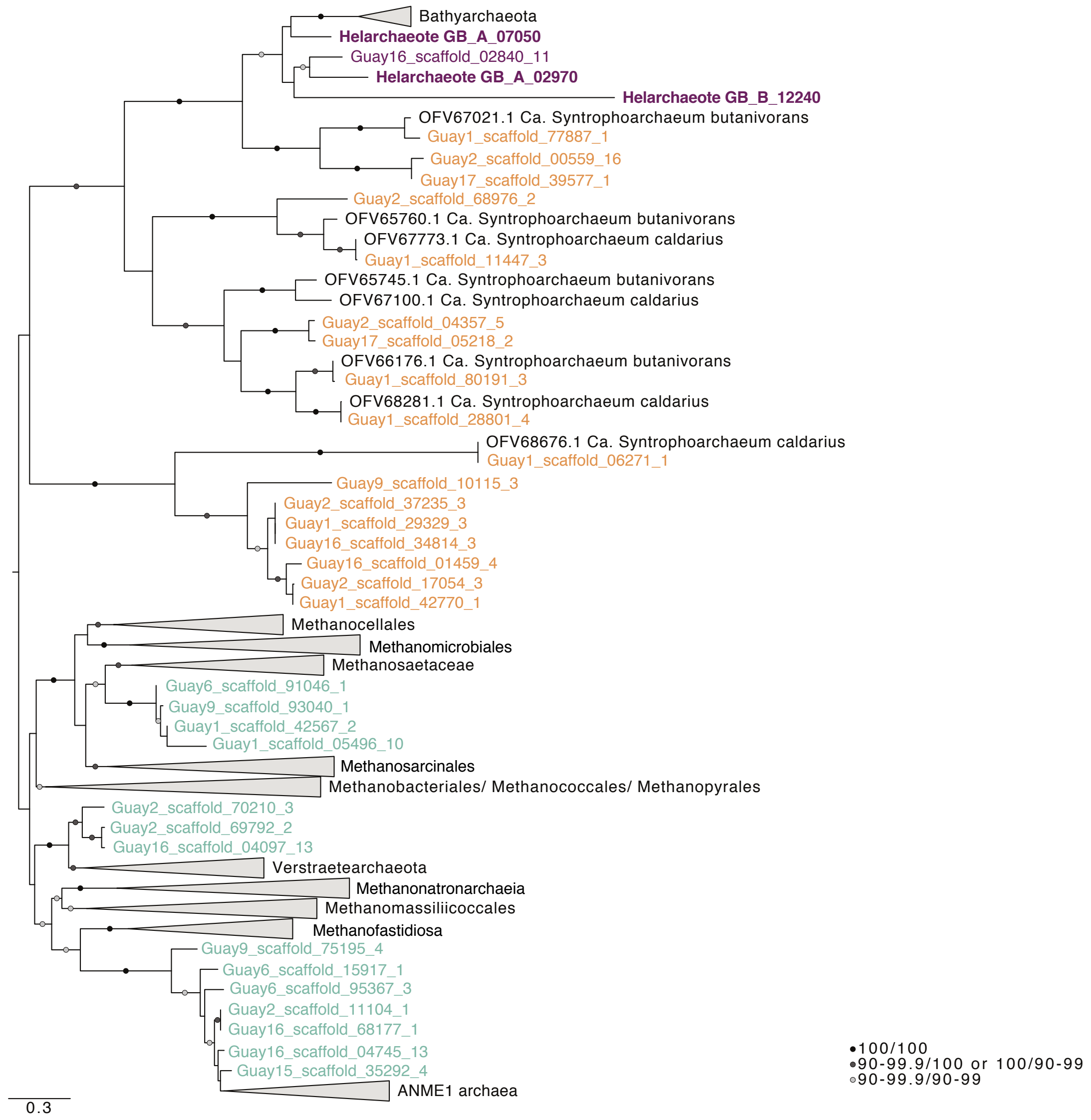


**Supplementary Figure 2.** Phylogenetic analysis of all McrA proteins recovered from unbinned GB scaffolds and Helarchaeota genomes as well as known homologs. Phylogenetic tree was inferred using Iq-tree (LG, C60, F, R) (See Supplementary Methods). Rapid and single branch test bootstraps above 90% are shown according to the color code given in the figure legend, i.e. black circles represent bootstrap values of 100/100, dark grey circles represent values of 90-99.9/100 and light grey represent values of 90-99.9/90-99. Helarchaeota McrA proteins group closest to McrA proteins identified in Bathyarchaeota spp. followed by those of *Ca*. Syntrophoarchaeum spp. Purple names represent Helarchaeota McrA, orange represents McrA versions proposed to use butane and green McrA types that are methane-specific. The tree was rooted arbitrarily between the cluster comprising canonical and divergent McrA homologs, respectively. Scale bars indicate the average number of substitutions per site.

| Identity | K256 | H257 | R271 | A272 | F330 | Y333 | Q400 | F443 | Y444 | G445 | C452 |
|---|---|---|---|---|---|---|---|---|---|---|---|
| Methanobrevibacter smithii ATCC 35061 YP_001273475.1 | **K** | **H** | **R** | **A** | **F** | **Y** | **Q** | **F** | **Y** | **G** | **C** |
| Candidatus Syntrophoarchaeum butanivorans OFV66176.1 | K | H | R | G | G | A | I | F | G | S | L |
| Candidatus Syntrophoarchaeum butanivorans OFV65745.1 | K | H | R | G | G | T | I | V | G | I | I |
| Candidatus Syntrophoarchaeum butanivorans OFV65760.1 | K | H | K | A | A | A | I | W | G | G | P |
| Candidatus Syntrophoarchaeum caldarius OFV67100.1 | K | H | R | G | G | T | I | V | G | V | L |
| Candidatus Syntrophoarchaeum caldarius OFV68676.1 | K | H | R | A | W | M | Q | F | Y | A | Q |
| Candidatus Syntrophoarchaeum caldarius OFV67773.1 | K | H | K | A | V | A | I | W | G | G | P |
| Bathyarchaeota CX-10 KT387810.1 | K | H | R | M | F | T | H | W | A | G | I |
| Bathyarchaeota BA2 KT387806.1 | K | H | R | M | F | V | H | W | A | G | I |
| Bathyarchaeota BA1 KT387805.1 | K | H | R | M | F | T | H | W | A | G | I |
| Hel_GB_A_02970 | K | S | R | A | F | V | H | W | A | G | L |
| Hel_GB_A_07050 | K | S | R | A | F | V | H | W | A | G | V |
| Hel_GB_B_12240 | Q | Y | L | G | I | V | Q | T | T | G | L |

**Supplementary Figure 3.** Comparison of amino acid active sites on the mcrA alignment. Numbers on top indicate position in alignment and expected amino acid. Letters correspond to amino acid one-letter code. Grey box shows reference genome Methanobrevibacter smithii ATCC 3506 and bold amino acid codes represent conserved residues for methanogenic archaea as previously described[29]. Grey letters represent conserved residues and black letter represents variation in the active sites.

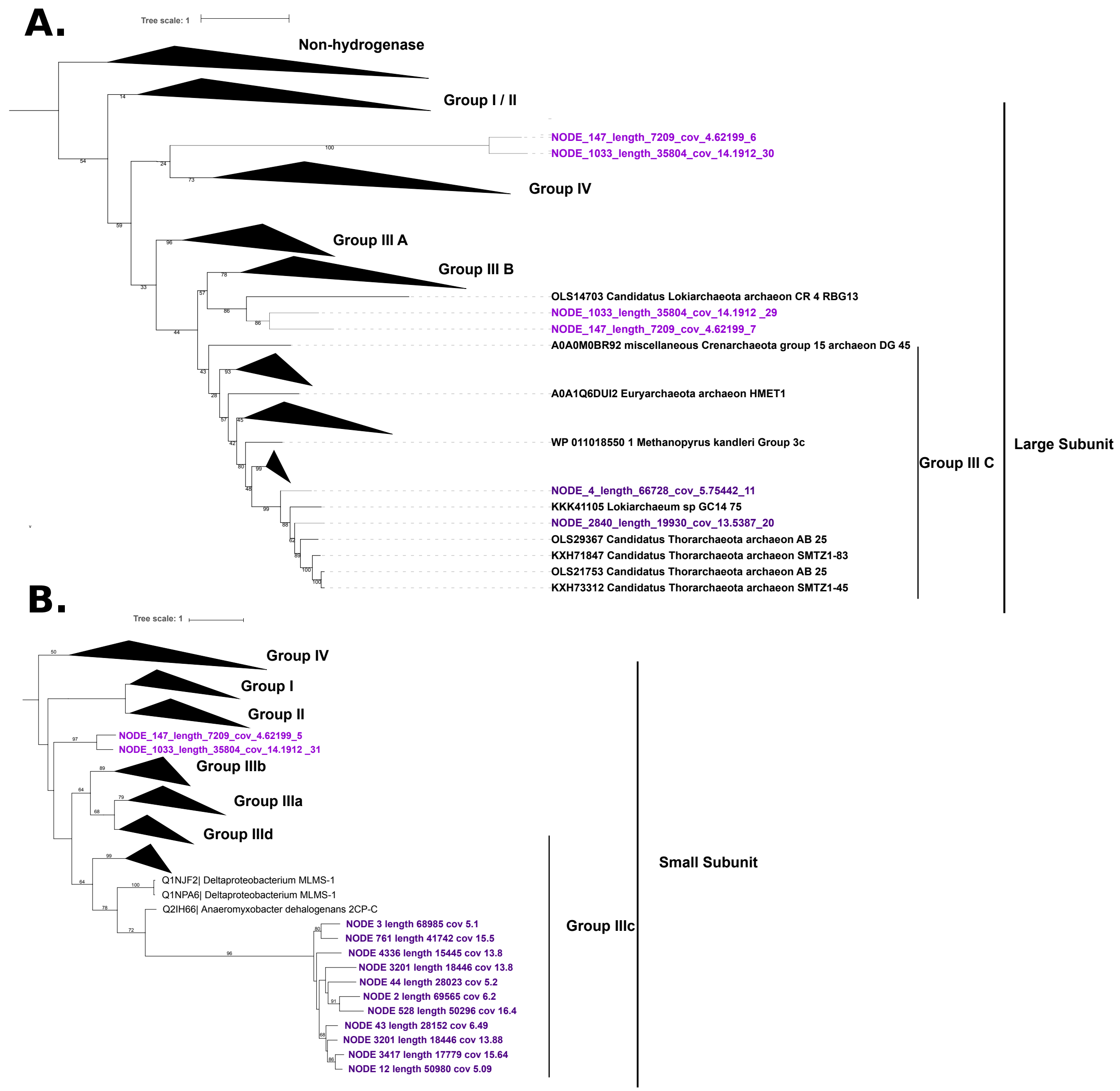


**Supplementary Figure 4.** Phylogenetic analyses of hydrogenases in Helarchaeota genomic bins. Both trees were generated using raxmlHPC-PTHREADS-AVX with GAMMA model parameters. (A) Large subunits of hydrogenases identified in Helarchaeota were aligned to a database[13,30] of well-characterized large subunits. Bootstrapping parameters were set to 100 generations. (B) Small subunits of hydrogenases identified in Helarchaeota were aligned to well-characterized small subunits[10,11] and raxml was run using the autoMRE setting resulting in 250 bootstrap generations.

### Supplementary Tables:

**Supplementary Table 1.** Geochemical and temperature data for sites from which genomes of Helarchaeota were assembled and binned.

| **SeqID** | **Hel_GB_A** | **Hel_GB_B** |
|---|---|---|
| Site | **4569_2** | **4571_4** |
| Sample | G16 | G9 |
| Beggiatoa Mat | Yes | No |
| Depth range | intermediate | shallow |
| Depth (cm) | 12-15 | 0-3 |
| $SO_4$ (mM) | 21.40 | 21.85 |
| Sulfide (mM) | 2.11 | 2.16 |
| $CH_{4\ (mM)}$ | 2.33 | 2.98 |
| δ13C of $CH_4$ | -27.62 | -34.32 |
| DIC (mM) | 16.58 | 10.20 |
| δ13/12 | -19.72 | -19.21 |
| Hydrothermal | yes | yes |
| Temperature range | medium | cool |
| T ($^o$C) | 28 | 10 |
| other | n/a | oily, sulfidic |

**Supplementary Table 2.** Percent identity for likely Helarchaeota 16S rRNA gene sequence compared to published Asgard and sequences identified in other proposed Helarchaeota bins[31].

| **Fields: query id** | **subject id** | **% identity** | **alignment length** | **mismatches** | **gap opens** | **q. start** | **q. end** | **s. start** | **s. end** | **evalue** | **bit score** |
|---|---|---|---|---|---|---|---|---|---|---|---|
| **Ga0180301_10078946** | Meg22_1214_Bin_228_scaffold_10652 | 96.5 | 400 | 13 | 1 | 3 | 401 | 1055 | 656 | 0 | 660 |
| **Ga0180301_10078946** | Meg22_1214_Bin_169_scaffold_22048 | 94.86 | 486 | 25 | 0 | 435 | 920 | 657 | 172 | 0 | 760 |
| **Ga0180301_10078946** | Odinarchaeota archaeon LCB_4 | 86.56 | 625 | 82 | 2 | 435 | 1058 | 922 | 1545 | 0 | 688 |
| **Ga0180301_10078946** | Lokiarchaeum sp. GC14_75 | 82.67 | 629 | 99 | 6 | 435 | 1058 | 867 | 1490 | 7.00E-159 | 549 |
| **Ga0180301_10078946** | Thorarchaeum sp SMTZ1-83 | 77.02 | 409 | 82 | 11 | 3 | 404 | 404 | 1 | 5.00E-61 | 224 |
| **Ga0180301_10078946** | Heimdallarchaeota archaeon AB_125 | 76.37 | 402 | 90 | 5 | 3 | 401 | 452 | 851 | 4.00E-57 | 211 |

**Supplementary Table 3.** Amino acid identity (AAI) comparison and estimated genome size between Helarchaeota bins and published Asgard genomes. AAI was performed by CompareM[32]. Estimated genome size for published Asgards from Zaremba-Niedzwiedzka, et al. 2017 (Supplementary Methods).

| **Genome A** | **Genes in A** | **Predicted Genome Size A (Mbp)** | **Genome B** | **Genes in B** | **Predicted Genome Size B (Mbp)** | **# orthologous genes** | **Mean AAI** | **Std AAI** | **Orthologous fraction (OF)** |
|---|---|---|---|---|---|---|---|---|---|
| Hel_GB_A | 3595 | 4.6 | Hel_GB_B | 3157 | 4.1 | 1477 | 51.96 | 12.63 | 46.78 |
| Hel_GB_A | 3595 | 4.6 | Odin.LCB.4_MDVT01000001.1 | 1580 | 1.5 | 574 | 45.89 | 9.94 | 36.33 |
| Hel_GB_A | 3595 | 4.6 | Loki.CR.4_MBAA01000194.1 | 4281 | 5.2 | 634 | 43.38 | 9.35 | 17.64 |
| Hel_GB_A | 3595 | 4.6 | Thor.AB.25_MEHG01000001.1 | 2763 | 3.0 | 553 | 43.02 | 8.91 | 20.01 |
| Hel_GB_A | 3595 | 4.6 | Heimdall.AB.125_MEHH01000001.1 | 2194 | 3.0 | 354 | 42.02 | 8.45 | 16.13 |
| Hel_GB_A | 3595 | 4.6 | Heimdall.LC3_MDVS01000001.1 | 5410 | 5.7 | 442 | 40.91 | 7.79 | 12.29 |
| Hel_GB_A | 3595 | 4.6 | Heimdall.LC2_MDVR01000001.1 | 4585 | 4.8 | 361 | 40.58 | 7.6 | 10.04 |
| Hel_GB_B | 3157 | 4.1 | Odin.LCB.4_MDVT01000001.1 | 1580 | 1.5 | 555 | 44.74 | 9.64 | 35.13 |
| Hel_GB_B | 3157 | 4.1 | Loki.CR.4_MBAA01000194.1 | 4281 | 5.2 | 624 | 43.15 | 9.15 | 19.77 |
| Hel_GB_B | 3157 | 4.1 | Thor.AB.25_MEHG01000001.1 | 2763 | 3.0 | 521 | 41.99 | 8.47 | 18.86 |
| Hel_GB_B | 3157 | 4.1 | Heimdall.AB.125_MEHH01000001.1 | 2194 | 3.0 | 359 | 41.43 | 7.95 | 16.36 |
| Hel_GB_B | 3157 | 4.1 | Heimdall.LC2_MDVR01000001.1 | 4585 | 4.8 | 362 | 40.51 | 7.58 | 11.47 |
| Hel_GB_B | 3157 | 4.1 | Heimdall.LC3_MDVS01000001.1 | 5410 | 5.7 | 441 | 40.31 | 7.52 | 13.97 |

**Supplementary Table 5.** Table of carbohydrate degrading enzymes identified in Helarchaeota bins.

| CAZy family | Defined activity | EC # or pfam | Hel_GB_A | Hel_GB_B |
|---|---|---|---|---|
| ***Cellulose degradation*** | | | | |
| GH5 | Cellulase | | 0 | 0 |
| | Endoglucanase | 3.2.1.4 | 0 | 0 |
| GH6 | Putative endoglucanases | | 0 | 0 |
| GH7 | | | 0 | 0 |
| GH9 | | | 0 | 0 |
| GH45 | | | 0 | 0 |
| GH48 | Endo-processive cellulases | | 0 | 0 |
| GH1 | β-glucosidase | | 0 | 0 |
| GH3 | | | 2 | 1 |
| **Total** | | | 2 | 1 |
| ***Hemicellulose -Debranching enzymes*** | | | | |
| GH51 | α-L-arabinofuranosidase | | 2 | 1 |
| GH54 | | | 0 | 0 |
| GH62 | | | 0 | 0 |
| | β-D-glucuronidase | 3.21.1.31 | 0 | 0 |
| GH67 | Putative α-D-glucuronidase | | 0 | 0 |
| | α-L-rhamnosidase | 3.2.1.40 | 0 | 0 |
| GH78 | putative rhamnosidase | | 1 | 1 |
| **Total** | | | 3 | 2 |
| ***Endohemicellases*** | | | | |
| GH53 | Endo-1,4-β-galactanase | | 0 | 0 |
| GH8 | Endo xylanase | | 0 | 0 |
| GH10 | Endo-1,4-β-xylanase | | 0 | 0 |
| GH11 | Xylanase | | 0 | 0 |
| GH28 | Putative galacturonases | | 0 | 0 |
| | Endo-mannanase | 3.2.1.78 | 0 | 0 |
| GH26 | Putative β mannanase and xylanase | | 0 | 0 |
| | α-D-xylosidase | 3.2.1._ | 0 | 0 |
| **Total** | | | 0 | 0 |
| ***Other Oligosaccharide-degrading enzymes*** | | | | |
| GH39 | β-1,4 -xylanase/β-D- | 3.2.1.37 | 0 | 0 |
| GH43 | Putative xylosidase and arbinases | | 0 | 0 |
| | β-1,4,-mannosidase | 3.2.1.25 | 0 | 0 |
| GH38 | α mannosidase | | 2 | 1 |
| GH2 | β-galactosidase | | 4 | 7 |
| GH35 | Putative β-galactosidase | | 0 | 1 |
| GH42 | | | 0 | 0 |
| GH53 | Endo-1,4-β-galactanase | | 0 | 0 |
| GH29 | α-L-fucosidase | | 2 | 0 |
| **Total** | | | 8 | 9 |
| ***Amylolytic enzymes*** | | | | |
| | α amylase | 3.2.1.1 | 0 | 0 |
| | gluco amylase | 3.2.1.3 | 0 | 0 |
| | pullanase | 3.2.1.41 | 0 | 0 |
| | isoamylase | 3.2.168 | 0 | 0 |
| **Total** | | | 0 | 0 |
| ***Other degradative enzymes*** | | | | |
| GH18 | Chitinase | | 0 | 0 |
| | β-N-acetyl-glucosaminidase (NAG) | 3.2.1.52 | 0 | 0 |
| | Pectin-esterase | 3.1.1.11 | 0 | 0 |
| | Pectin lyase | 4.2.2.10 | 0 | 0 |
| | Putative pectin hydrolysis | PF01095, PF00544 | 0 | 0 |
| **Total** | | | 0 | 0 |

**Supplementary Table 6.** List of 37 Phylosift[33] marker genes that were used for preliminary phylogenetic identification of individual bins

| PhyloSift Marker | Gene Name |
|---|---|
| DNGNGWU00001 | ribosomal protein S2 rpsB |
| DNGNGWU00002 | ribosomal protein S10 rpsJ |
| DNGNGWU00003 | ribosomal protein L1 rplA |
| DNGNGWU00005 | translation initiation factor IF-2 |
| DNGNGWU00006 | metalloendopeptidase |
| DNGNGWU00007 | ribosomal protein L22 |
| DNGNGWU00009 | ribosomal protein L4/L1e rplD |
| DNGNGWU00010 | ribosomal protein L2 rplB |
| DNGNGWU00011 | ribosomal protein S9 rpsI |
| DNGNGWU00012 | ribosomal protein L3 rplC |
| DNGNGWU00013 | phenylalanyl-tRNA synthetase beta subunit |
| DNGNGWU00014 | ribosomal protein L14b/L23e rplN |
| DNGNGWU00015 | ribosomal protein S5 |
| DNGNGWU00016 | ribosomal protein S19 rpsS |
| DNGNGWU00017 | ribosomal protein S7 |
| DNGNGWU00018 | ribosomal protein L16/L10E rplP |
| DNGNGWU00019 | ribosomal protein S13 rpsM |
| DNGNGWU00020 | phenylalanyl-tRNA synthetase alpha subunit |
| DNGNGWU00021 | ribosomal protein L15 |
| DNGNGWU00022 | ribosomal protein L25/L23 |
| DNGNGWU00023 | ribosomal protein L6 rplF |
| DNGNGWU00024 | ribosomal protein L11 rplK |
| DNGNGWU00025 | ribosomal protein L5 rplE |
| DNGNGWU00026 | ribosomal protein S12/S23 |
| DNGNGWU00027 | ribosomal protein L29 |
| DNGNGWU00028 | ribosomal protein S3 rpsC |
| DNGNGWU00029 | ribosomal protein S11 rpsK |
| DNGNGWU00030 | ribosomal protein L10 |
| DNGNGWU00031 | ribosomal protein S8 |
| DNGNGWU00032 | tRNA pseudouridine synthase B |
| DNGNGWU00033 | ribosomal protein L18P/L5E |
| DNGNGWU00034 | ribosomal protein S15P/S13e |
| DNGNGWU00035 | Porphobilinogen deaminase |
| DNGNGWU00036 | ribosomal protein S17 |
| DNGNGWU00037 | ribosomal protein L13 rplM |
| DNGNGWU00039 | ribonuclease HII |
| DNGNGWU00040 | ribosomal protein L24 |

**Supplementary Table 7.** List of selected phylogenetic markers used to reconstruct the organismal phylogeny (Fig. 1).

| |
|---|
| RP-L15; large subunit ribosomal protein L15 |
| RP-L18e; large subunit ribosomal protein L18e |
| RP-L32e; large subunit ribosomal protein L32e |
| RP-S14; small subunit ribosomal protein S14 |
| RP-L29; large subunit ribosomal protein L29 |
| RP-S19e; small subunit ribosomal protein S19e |
| RP-S13; small subunit ribosomal protein S13 |
| RP-S10; small subunit ribosomal protein S10 |
| RP-S17e; small subunit ribosomal protein S17e |
| RP-S6e; small subunit ribosomal protein S6e |
| RP-L24e; large subunit ribosomal protein L24e |
| RP-L40e; large subunit ribosomal protein L40e |
| RP-L2; large subunit ribosomal protein L2 |
| RP-L3; large subunit ribosomal protein L3 |
| RP-L4e; large subunit ribosomal protein L4e |
| RP-L23; large subunit ribosomal protein L23 |
| RP-L30; large subunit ribosomal protein L30 |
| RP-S5; small subunit ribosomal protein S5 |
| RP-L18; large subunit ribosomal protein L18 |
| RP-L19e; large subunit ribosomal protein L19e |
| RP-L6; large subunit ribosomal protein L6 |
| RP-S8; small subunit ribosomal protein S8 |
| RP-L5; large subunit ribosomal protein L5 |
| RP-S4e; small subunit ribosomal protein S4e |
| RP-L24; large subunit ribosomal protein L24 |
| RP-L14; large subunit ribosomal protein L14 |
| RP-S17; small subunit ribosomal protein S17 |
| RP-S3; small subunit ribosomal protein S3 |
| RP-L22; large subunit ribosomal protein L22 |
| RP-S19; small subunit ribosomal protein S19 |
| RP-S27e; small subunit ribosomal protein S27e |
| RP-L44e; large subunit ribosomal protein L44e |
| RP-L10e; large subunit ribosomal protein L10e |
| RP-L37e; large subunit ribosomal protein L37e |
| RP-L21e; large subunit ribosomal protein L21e |
| RP-S8e; small subunit ribosomal protein S8e |
| RP-L39e; large subunit ribosomal protein L39e |
| RP-S24e; small subunit ribosomal protein S24e |
| RP-S27Ae; small subunit ribosomal protein S27Ae |
| RP-S15; small subunit ribosomal protein S15 |
| RP-S3Ae; small subunit ribosomal protein S3Ae |
| RP-L37Ae; large subunit ribosomal protein L37Ae |
| RP-L15e; large subunit ribosomal protein L15e |
| RP-S4; small subunit ribosomal protein S4 |
| RP-S11; small subunit ribosomal protein S11 |
| RP-L13; large subunit ribosomal protein L13 |
| RP-S9; small subunit ribosomal protein S9 |
| RP-S2; small subunit ribosomal protein S2 |
| RP-S7; small subunit ribosomal protein S7 |
| RP-S12; small subunit ribosomal protein S12 |
| RP-L12; large subunit ribosomal protein L12 |
| RP-L10; large subunit ribosomal protein L10 |
| RP-L1; large subunit ribosomal protein L1 |
| RP-S28e; small subunit ribosomal protein S28e |
| RP-L11; large subunit ribosomal protein L11 |
| RP-L31e; large subunit ribosomal protein L31e |

**Supplementary Table 8.** Summary of phylogenetic analyses of 56 concatenated ribosomal proteins. Left column correspond to datasets varying in taxon sampling ("full", i.e., initial dataset, "without DPANN", "without Eukaryotes", "without DPANN and eukaryotes") or with unchanged taxon sampling but fastest-evolving sites removed (Full-FSR). Some or all of those have been subjected to amino-acid recoding (second column) and phylopgenetic reconstruction in a Maximum Likelihood or Bayesian framework (first column), under various models of evolution (third column). In each case, the first and second value correspond to the statistical support (PMSF boostrap or Posterior Probability) for the monophyly of Helarchaetoa and Lokiarchaeota, and of eukaryotes and Heimdallarchaeota, respectively. In addition, for Phylobayes analyses, are indicated the number of generations run, and the maxdiff between chains indicating convergence. An asterisk indicates that converge was obtained for three chains out of four only.

| | Iqtree | Phylobayes | |
|---|---|---|---|
| | Non-recoded | | SR4 recoding |
| | LG+C60+F+G+PMSF | CAT+LG | CAT+GTR |
| **Full** | 91; 82 | NA | 0.99; 0.76; 49305; 0.19 |
| **Full-FSR** | 95; 90 | NA | NA |
| **Without DPANN** | 89; 90 | 1.0; 0.99; 37831; 0.29* | NA |
| **Without Eukaryotes** | 91; - | NA | NA |
| **Without DPANN and Euk** | 93; - | NA | NA |

### Supplementary References:


1. Dombrowski, N., Seitz, K. W., Teske, A. P. & Baker, B. J. Genomic insights into potential interdependencies in microbial hydrocarbon and nutrient cycling in hydrothermal sediments. *Microbiome* **5**, 106 (2017).
2. Dick, G. J. *et al.* Community-wide analysis of microbial genome sequence signatures. *Genome Biology* **10**, R85 (2009).
3. Kang, D. D., Froula, J., Egan, R. & Wang, Z. MetaBAT, an efficient tool for accurately reconstructing single genomes from complex microbial communities. *PeerJ* **3**, e1165 (2015).
4. Alneberg, J. *et al.* Binning metagenomic contigs by coverage and composition. *Nature Methods* **11**, nmeth.3103 (2014).
5. Eren, A. M. *et al.* Anvi'o: an advanced analysis and visualization platform for 'omics data. *PeerJ* **3**, e1319 (2015).

6. Li, H. Aligning sequence reads, clone sequences and assembly contigs with BWA-MEM. *arXiv:1303.3997 [q-bio]* (2013).
7. Li, H. *et al.* The Sequence Alignment/Map format and SAMtools. *Bioinformatics* **25**, 2078–2079 (2009).
8. Sieber, C. M. K. *et al.* Recovery of genomes from metagenomes via a dereplication, aggregation, and scoring strategy. *bioRxiv* 107789 (2017). doi:10.1101/107789
9. Dombrowski, N., Teske, A. P. & Baker, B. J. Extensive metabolic versatility and redundancy in microbially diverse, dynamic Guaymas Basin hydrothermal sediments. *Nature Communications* **9:4999**, (2018).
10. Vignais, P. M. & Billoud, B. Occurrence, classification, and biological function of hydrogenases: an overview. *Chem. Rev.* **107**, 4206–4272 (2007).
11. Vignais, P. M., Billoud, B. & Meyer, J. Classification and phylogeny of hydrogenases1. *FEMS Microbiology Reviews* **25**, 455–501
12. Kearse, M. *et al.* Geneious Basic: an integrated and extendable desktop software platform for the organization and analysis of sequence data. *Bioinformatics* **28**, 1647–1649 (2012).
13. Spang, A. *et al.* A renewed syntrophy hypothesis for the origin of the eukaryotic cell based on comparative analysis of Asgard archaeal metabolism. *Nature Microbiology* **Submitted**,
14. Jones, P. *et al.* InterProScan 5: genome-scale protein function classification. *Bioinformatics* **30**, 1236–1240 (2014).

15. Krogh, A., Larsson, B., von Heijne, G. & Sonnhammer, E. L. Predicting transmembrane protein topology with a hidden Markov model: application to complete genomes. *J. Mol. Biol.* **305**, 567–580 (2001).

16. Pasquier, C., Promponas, V. J., Palaios, G. A., Hamodrakas, J. S. & Hamodrakas, S. J. A novel method for predicting transmembrane segments in proteins based on a statistical analysis of the SwissProt database: the PRED-TMR algorithm. *Protein Eng.* **12**, 381–385 (1999).

17. Käll, L., Krogh, A. & Sonnhammer, E. L. L. A combined transmembrane topology and signal peptide prediction method. *J. Mol. Biol.* **338**, 1027–1036 (2004).

18. Edgar, R. C. MUSCLE: multiple sequence alignment with high accuracy and high throughput. *Nucleic Acids Res* **32**, 1792–1797 (2004).

19. Criscuolo, A. & Gribaldo, S. BMGE (Block Mapping and Gathering with Entropy): a new software for selection of phylogenetic informative regions from multiple sequence alignments. *BMC evolutionary biology* **10**, 210 (2010).

20. Nguyen, L.-T., Schmidt, H. A., von Haeseler, A. & Minh, B. Q. Iq-tree: A fast and effective stochastic algorithm for estimating maximum-likelihood phylogenies. *Molecular biology and evolution* **32**, 268–274 (2015).

21. Finn, R. D. *et al.* The Pfam protein families database. *Nucleic Acids Res* **38**, D211–D222 (2010).

22. Söding, J., Biegert, A. & Lupas, A. N. The HHpred interactive server for protein homology detection and structure prediction. *Nucleic Acids Res* **33**, W244–W248 (2005).

23. Letunic, I., Doerks, T. & Bork, P. SMART: recent updates, new developments and status in 2015. *Nucleic Acids Res.* **43**, D257-260 (2015).

24. Klinger, C. M., Spang, A., Dacks, J. B. & Ettema, T. J. G. Tracing the Archaeal Origins of Eukaryotic Membrane-Trafficking System Building Blocks. *Mol. Biol. Evol.* **33**, 1528–1541 (2016).

25. Katoh, K. & Standley, D. M. MAFFT multiple sequence alignment software version 7: Improvements in performance and usability. *Molecular Biology and Evolution* **30**, 772–780 (2013).

26. Capella-Gutiérrez, S., Silla-Martínez, J. M. & Gabaldón, T. trimAl: a tool for automated alignment trimming in large-scale phylogenetic analyses. *Bioinformatics* **25**, 1972–1973 (2009).

27. Price, M. N., Dehal, P. S. & Arkin, A. P. FastTree: Computing Large Minimum Evolution Trees with Profiles instead of a Distance Matrix. *Mol Biol Evol* **26**, 1641–1650 (2009).

28. Hyatt, D. *et al.* Prodigal: prokaryotic gene recognition and translation initiation site identification. *BMC Bioinformatics* **11**, 119 (2010).

29. Hallam, S. J., Girguis, P. R., Preston, C. M., Richardson, P. M. & DeLong, E. F. Identification of Methyl Coenzyme M Reductase A (mcrA) Genes Associated with Methane-Oxidizing Archaea. *Appl Environ Microbiol* **69**, 5483–5491 (2003).

30. Søndergaard, D., Pedersen, C. N. S. & Greening, C. HydDB: A web tool for hydrogenase classification and analysis. *Sci Rep* **6**, 34212 (2016).

31. Altschul, S. F. *et al.* Gapped BLAST and PSI-BLAST: a new generation of protein database search programs. *Nucleic Acids Res.* **25**, 3389–3402 (1997).

32. https://github.com/dparks1134/CompareM.

33. Darling, A. E. *et al.* PhyloSift: phylogenetic analysis of genomes and metagenomes. *PeerJ* **2**, e243 (2014).
